# Reproductive interference hampers species coexistence despite conspecific sperm precedence

**DOI:** 10.1101/287482

**Authors:** Ryosuke Iritani, Suzuki Noriyuki

**Author notes:** Both authors contributed equally.

## Abstract

Negative interspecific mating interactions, known as reproductive interference, can hamper species coexistence in a local patch and promote niche partitioning or geographical segregation of closely related species. Conspecific sperm precedence (CSP), which occurs when females that have mated with both conspecific and heterospecific males preferentially use conspecific sperm for fertilization, might contribute to species coexistence by mitigating the costs of interspecific mating and hybridization. We examined whether two closely related species exhibiting CSP can coexist in a local environment in the presence of reproductive interference. First, using a behaviourally explicit mathematical model, we demonstrated that two species characterized by negative mating interactions are unlikely to coexist because the costs of reproductive interference, such as loss of mating opportunity with conspecific partners, are inevitably incurred when individuals of both species are present. Second, we experimentally demonstrated differences in mating activity and preference in two *Harmonia* ladybird species known to exhibit CSP. According to the developed mathematical model of reproductive interference, these behavioural differences should lead to local extinction of *H. yedoensis* because of reproductive interference by *H. axyridis*. This prediction is consistent with field observations that *H. axyridis* uses various food sources and habitats whereas *H. yedoensis* is confined to a less preferred prey item and a pine tree habitat. Finally, by a comparative approach, we showed that niche partitioning or parapatric distribution, but not sympatric coexistence in the same habitat, is maintained between species with CSP belonging to a wide range of taxa, including vertebrates and invertebrates living in aquatic or terrestrial environments. Taken together, these results lead us to conclude that reproductive interference generally destabilizes local coexistence even in closely related species that exhibit CSP.

## Introduction

Restrictions to local coexistence among phylogenetically related species are closely related to niche partitioning and the diversification of resource use traits, which help to determine community assemblages at both local and regional scales (Schluter 2000, Grant and Grant 2011, Losos 2011). Therefore, understanding the mechanisms that restrict local coexistence is of fundamental importance in ecology and evolution. Negative interspecific mating interaction, that is, reproductive interference, is one mechanism that can drive species exclusion at local scale and subsequent niche partitioning among species (Gröning and Hochkirch 2008). Reproductive interference has been theoretically demonstrated to hamper species coexistence in a homogeneous environment even in ecologically neutral species with similar growth rates and abilities to compete for shared resources (Kuno 1992, Konuma and Chiba 2007, Crowder et al. 2011, Nishida et al. 2015, Kyogoku and Sota 2017). Moreover, empirical studies have also reported that reproductive interference contributes to niche partitioning between congeneric species with overlapping mating signals, including in frogs (Ficetola and Bernardi 2005), birds (Vallin et al. 2012), mites (Takafuji et al. 1997), and insects (butterflies, Friberg et al. 2013; grasshoppers, Hochkirch et al. 2007; ladybirds, Noriyuki et al. 2012). Therefore, reproductive interference is a determinant of local and regional species diversity in a wide range of animal taxa in nature, though its significance in community ecology has been underestimated for decades (Gröning and Hochkirch 2008, Kyogoku 2015).

A number of mechanisms, however, are reported to mitigate the negative impacts of reproductive interference on the coexistence of species occupying the same niche, including plastic responses in reproductive traits (Otte and Hilker 2016), continued dispersal to new sets of ephemeral resource patches (Ruokolainen and Hanski 2016), and reinforcement of reproductive isolation (Bargielowski et al. 2013). One possible mitigating mechanism is conspecific sperm precedence (CSP), where females that have mated with both conspecific and heterospecific males preferentially use conspecific sperm for fertilization (Howard 1999). Such females might experience fewer costs associated with interspecific mating and hybridization (i.e., waste of gametes), because most or all of their offspring will be pure conspecifics (Nakano 1985, Veen et al. 2001, Marshall et al. 2002). In addition, in various animals, mating order has been shown to have no influence on whether a female us able to preferentially use conspecific sperm (Howard et al. 1998, Marshall et al. 2002), suggesting that complete CSP can largely eliminate the negative impact of interspecific mating provided that females have mated with at least one conspecific male before the onset of oviposition or birthing (Marshall et al. 2002). CSP has been reported in a variety of animal taxa, including sea urchins (Geyer and Palumbi 2005), mussels (Klibansky and McCartney 2013), crickets (Howard et al. 1998), fruit flies (Price 1997), beetles (Fricke and Arnqvist 2004, Rugman-Jones and Eady 2007), fishes (Yeates et al. 2013), and mice (Dean and Nachman 2009), and thus potentially plays an important role in species coexistence. Although CSP has attracted much attention as a driver of speciation through post-mating and pre-zygotic reproductive isolation (Howard et al. 1998; Howard 1999), it is still unclear whether CSP can sufficiently ameliorate the cost of reproductive interference to promote stable coexistence of closely related species in the same local environment.

CSP may not fully function as a barrier against reproductive interference. Under imperfect species discrimination, individual females may incur a variety of costs as a result of interactions with heterospecific males during the reproductive process, such as reduced longevity and oviposition rates (Kawatsu and Kishi 2017), physical damage caused by interspecific copulation (Kyogoku and Sota 2015), and loss of opportunity to mate with conspecific partners (Thum 2007, Noriyuki et al. 2012, Ramiro et al. 2015), as well as the production of unviable hybrid offspring (Todesco et al. 2016). CSP alone might be insufficient to compensate all of these potential costs of reproductive interference. In addition, adaptive behaviours of females and males can prevent multiple matings by females and consequently make the CSP mechanism useless. For example, studies on sexual conflict have shown that females are likely to avoid multiple matings when the benefit is low (Eberhard 1996, Arnqvist and Rowe 2005). Moreover, to prevent sperm competition, males often try to prevent females from mating multiple times, for example, by mate guarding after copulation (Alcock 1994), by placing a physical plug in female reproductive organs (Matsumoto and Suzuki 1992, Polak et al. 2001), or by insertion of a chemical that inhibits remating receptivity (Scott 1986, Gillott 2003, Himuro and Fujisaki 2008). Therefore, to evaluate the ecological role of CSP in species coexistence, various behavioural and physiological mechanisms affecting the reproductive process must be taken into account.

In this study, we examined whether CSP can mitigate the effect of reproductive interference in two closely related species so that they are able to coexist in a local environment. We adopted a tripartite approach. First, we developed a behaviourally explicit mathematical model to analyse behavioural and demographic factors affecting local species coexistence, with a focus on the multiple copulation rate, mating preference toward conspecific or heterospecific partners, and the initial population densities of the two species. Second, we conducted mating experiments with two predatory ladybird species, *Harmonia axyridis* and *Harmonia yedoensis*, to test the predictions of the mathematical model. CSP has been detected in both these species (Noriyuki et al. 2012), and they occupy different niches in nature; *H. axyridis* is a generalist that feeds on various species of preferred aphids, whereas *H. yedoensis* specializes on the giant pine aphid, which is a highly elusive prey item and nutritionally poor for larval development (Noriyuki et al. 2011, Noriyuki and Osawa 2012). In addition, the reproductive success of *H. yedoensis* females is strongly decreased in the presence of *H. axyridis* males, suggesting that *H. yedoensis* might utilize the less preferred food and habitat to avoid reproductive interference from *H. axyridis* (Noriyuki et al. 2012). Third, we investigated the general consequences of CSP on species coexistence in nature by compiling published data on pairs of species in which CSP has been detected and found that such species pairs generally show niche separation (habitat and food source) or geographically separate distributions. We concluded from our results that CSP does not reduce the overall cost of reproductive interference sufficiently to allow the interacting species to coexist in the same local environment.

## Materials and methods

### Mathematical model

We modelled a community of two species (X and Y), with density *N*_X_(*t*) and *N*_Y_(*t*), respectively, in generation $, inhabiting a single patch. The two species interact through resource competition as well as through reproductive interference, but they are ecologically neutral in terms of the total number of offspring per capita that survive to maturation (denoted *r*), density-dependent regulation (denoted ’), and interspecific competitive strength (denoted *b*). We assumed a sex ratio of 1:1 (though we found that the ratio does not affect the results; see Kyogoku and Sota 2017), and, for the sake of simplicity, at most two instances of copulation per female. Finally, we assumed that females are not always capable of correctly assessing the species identity of their mating partner; as a result, interspecific mating can occur even after intraspecific mating (as is the case in *H. yedoensis* and *H. axyridis*).

Species X and Y can differ with respect to the rate at which females accept males as mates (Fig. 1). Specifically, a virgin X-female (i.e., a female of species X) accepts a mating attempt by an X-male with probability *p*_X|X_ and a Y-male with probability *p*_X|Y_, and a once-mated female accepts a mating attempt by an X-male with probability *q*_X|X_ and with a Y-male with probability *q*_X|Y_. Similarly, the probabilities of a Y-female accepting a mating attempt by a male in the corresponding situations are *p*_Y|Y_, *p*_Y|X_, *q*_Y|Y_ and *q*_Y|X_.

**Fig. 1.**
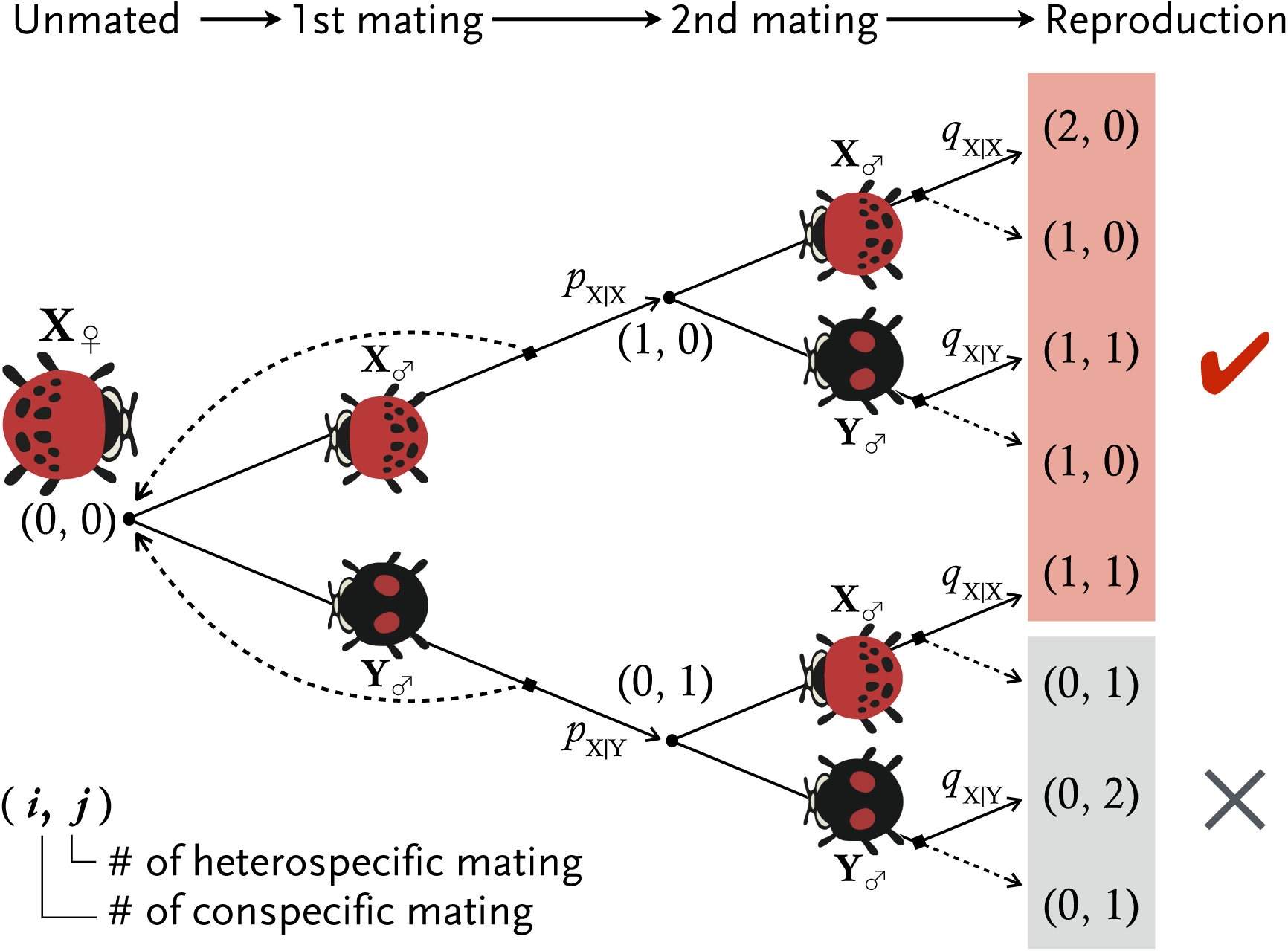
Schematic mating decision-making tree for a female of species X according to our mathematical model. Here, (P, Q) means a female with *i* intraspecific matings and *j* interspecific matings (1 ≤ *i* + *j* ≤ 2). A virgin female has state (0, 0), and she accepts a given X-male or Y-male with a probability *p*_X|X_ and *p*_X|Y_, respectively. Subsequently, the non-virgin female with state (0, 1) or (1, 0) accepts an X-male or Y-male with probability *q*_X|X_ or *q*_X|Y_, respectively. The corresponding mating decision-making tree for a Y-female can be obtained by exchanging X and Y. The female states after the second mating that include at least one intraspecific mating (*i* ≥ 1) are shaded red; in this case, the female can produce offspring of her own species through CSP. The states of females that failed to copulate with a conspecific male before producing offspring (*i* = 0) are shaded grey.

The frequencies of X-males and Y-males (among all males in the community) are as follows:

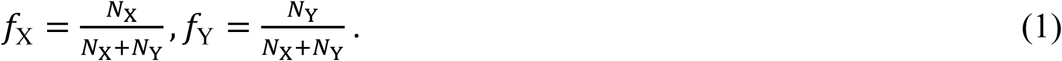

We denote the expected reproductive output by a single X-female or a single Y-female by *E*_X_or *E*_Y_, respectively. Parameter 4 tunes the intensity of reproductive interference (0 < 4 < 1) and reflects the degree of interspecific overlap in the reproductive niche; thus, the expected reproductive output of an X- or Y-female is calculated as:

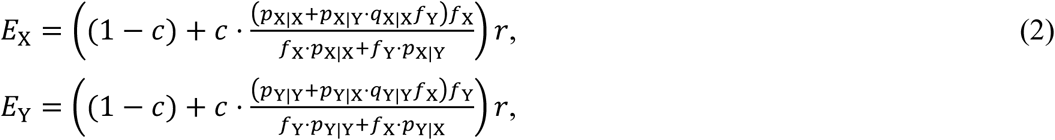

(see Appendix A). Within the parentheses on the right side of each Eq. (2), (1 − 4) represents reproductive success independent of density and frequency, and the second term represents the product of reproductive interference intensity (*c*) and the conditional probability that, given a non-virgin, a single female mates with a conspecific male at least once.

To model the population dynamics under intra- and interspecific competition, we used the Beverton–Holt model of community dynamics (Beverton and Holt 1957, May and Oster 1976, Ackleh and Salceanu 2014). Specifically, we assume that regulation occurs among adults, followed by reproduction. Under this assumption, the dynamics are as follows:

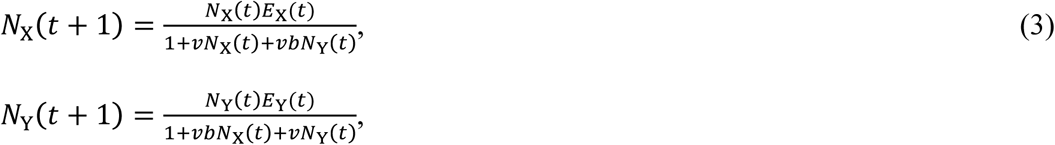

where $ represents the generation, > ≥ 1 represents the life-time survival rate (subsuming the total number of eggs per capita), ’ ≥ 0 tunes density dependence in regulation, and *b* (0 ≤ F ≤ 1) tunes the strength of interspecific resource competition. Throughout this analysis, we set ’ = 1, which does not cause any loss of generality (Ackleh and Salceanu 2014). By using the underlying link between a continuous-time logistic equation and the discrete-time Beverton–Holt model (May and Oster 1976), we approximate the dynamics by the following ordinary differential equations (ODE):

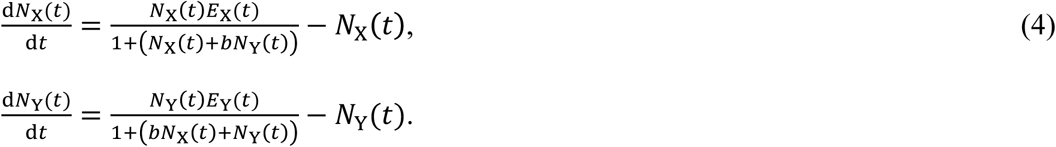

All variables and parameters are defined in Table 1. The community equilibrium is obtained by setting Eqs. (4) to zero. We also carry out a basic local stability analysis of the equilibrium of the dynamical system to determine possible equilibrium states. Specifically, we identified conditions leading to *species exclusion* (i.e., only one species persists) or *coexistence* (i.e., both species coexist).

**Table 1.** Parameters included in the model.

| Parameter/Variable | Definition | Default value (if any) |
| --- | --- | --- |
| X or Y | Species label | – |
| $N_X, N_Y$ | Density of X or Y | Dynamic variable |
| $t$ | Generation ($t \geq 0$) | – |
| $r$ | Egg production (per capita) | 25 |
| $v$ | Density dependence of resource competition ($v > 0$) | 1 |
| $b$ | Strength of interspecific competition ($b > 0$) | 0.3 |
| $c$ | Strength of reproductive interference ($0 < c < 1$); probability that a female is subject to a possible heterospecific mating | Varied |
| $p_{i\vert j}$ | Probability that a virgin female of species $i$ accepts a mating attempt by a male of species $j$, where $i$ and $j$ can be either X or Y | See Fig. 2 |
| $q_{i\vert j}$ | Probability that a non-virgin female of species $i$ accepts a mating attempt by a male of species $j$, where $i$ and $j$ can be either X or Y | See Fig. 2 |
| $f_X (= N_X/(N_X + N_Y))$ | Frequency of species X in the population | Dynamic variable |
| $E_X, E_Y$ | Expected reproductive output (per capita), calculated based upon the mating decision-making tree (see Fig. 1), for species X or Y | – |

We also visualized the steady states by a numerical approach, first (i) evaluating the eigenvalues of the Jacobi matrix of equilibria and then (ii) depicting the phase portraits (using Mathematica 11.2.0; Wolfram Research 2017). For the eigenvalue analyses, we first checked the number of feasible equilibria (*N*_X_, *N*_Y_ ≥ 0) given the community dynamics and then numerically evaluated the real part of the eigenvalues associated with the corresponding equilibria.

### Experiment

We collected adults of two ladybird species from Japanese red pine (*Pinus densiflora* Sieb. et Zucc.) at the University of Tokyo Tanashi Forest (139°32’E, 35°44’N), Tanashi city, Tokyo, during April 2014, and at the Kumagaya campus of Rissho University (139°36’E, 36°10’N) and the Hirose Wild Birds Forest (139°35’E, 36°14’N), Kumagaya city, central Japan, during April 2015. In the laboratory, we maintained females individually in plastic Petri dishes (9 cm in diameter by 1.5 cm high) at 25 °C, and fed them each day with a surplus of frozen *Ephestia kuehniella* Zeller eggs (Beneficial Insectary, Ontario, Canada) to ready them for reproduction. In total, 15 *H. yedoensis* and 8 *H. axyridis* females in 2014 and 10 *H. yedoensis* and 9 *H. axyridis* females in 2015 produced a sufficient number of egg clutches for our experiments. In addition, in 2015, we collected 32 *H. yedoensis* egg clutches and 41 *H. axyridis* egg clutches that had been oviposited on the leaves and branches of Japanese red pine trees at the Hirose Wild Birds Forest. We fed the hatched offspring from both laboratory-laid and wild-collected egg clutches with a mixture of sucrose, dried yeast, and powdered drone honeybee (following Niijima et al. 2000) to the adult stage in plastic cases (each 12.5 cm in diameter by 9.5 cm high) containing wood wool as a substrate on which they could walk. We recorded the date of emergence, body length (to the nearest 0.01 mm), and elytra colour (black or red) of all newly emerged adults as possible factors affecting mating preference, and used these virgin individuals for the following behavioural experiments to standardize the mating experience.

Because it takes approximately 1 month for most individuals of both *H. yedoensis* and *H. axyridis* to mature sexually after they emerge as adults (Okada, Nijima & Toriumi 1978), we reared the newly emerged adults individually in plastic Petri dishes for at least 30 days, providing them with frozen *E. kuehniella* eggs every other day, before using them in mating experiments. In addition, we excluded egg clutches from the wild-caught mothers that produced only female offspring (two *H. yedoensis* females in 2014 and one *H. axyridis* female in 2015) because they were likely to be infected with male-killing bacteria (Noriyuki et al. 2014, 2016), to avoid any confounding effects of male-killing bacteria on the host mating behaviour (Majerus 2003).

In the mating experiment, we kept one female (*H. yedoensis* or *H. axyridis*) and one male (*H. yedoensis* or *H. axyridis*) together in a small Petri dish (5 cm in diameter) on a laboratory bench at room temperature (25 °C) under constant fluorescent lighting. We never placed females with sibling males (i.e., individuals produced by the same wild-caught mother or from the same wild-collected clutch) to preclude any effects of inbreeding avoidance on mating behaviour. We observed the occurrence of male mating attempts, female rejection behaviour, and successful copulation in each experimental session (see Noriyuki et al. 2012 for the definition of these behaviours). In 2014, we visually observed mating activities during 15-min sessions. In 2015, we used videocameras (HC-V480, Panasonic, Osaka, Japan) to record experimental sessions for at least 6 hours (up to 20 hours) and then watched the videos to analyse mating behaviours. In the 2014 experiments, each pair was allowed to mate after the 15-min session until copulation was completed. In the 2015 experiments, multiple copulations were allowed in the same experimental session. Note that Noriyuki et al. (2012) reported that the mean duration of copulation was 228 min in *H. yedoensis* and 124 min in *H. axyridis* under similar experimental conditions. In both 2014 and 2015, we reused virgin and non-virgin individuals after the experimental session for other sessions to analyse the effects of mating experience on subsequent mating behaviour.

To examine the effect of mating experience in virgins and non-virgins on the copulation rate in each species, we analysed the proportion of experimental sessions that included successful copulation (at least one in the 2015 experiments) by a generalized linear mixed model with a binomial error structure using the glmer function of the lme4 library (Bates et al 2015) of the R software package (version 3.4.2, R Core Team 2017). Similarly, we compared the mating rate between intra- and interspecific mating trials in virgin and non-virgin females. Moreover, we analysed mating preferences of both males and females to determine factors responsible for the copulation rate. First, we evaluated male preference by the proportion of experimental sessions that included at least one male mating attempt, whether or not it was followed by successful mating. Second, we examined the female preference by calculating the proportion of male mating attempts that elicited female rejection behaviour. In all analyses, we also incorporated the date of emergence, body length, and elytra colour of females and males as fixed effects, and the identity of the mother of the female and that of the male as a random term. We analysed data from the experiments in 2014 and 2015 separately because of the differences in the source populations and the specific experimental conditions.

Furthermore, we applied signal detection theory (Green and Swets 1966) to disentangle the mechanism of decision making in males and females who need to choose conspecific mating partner over heterospecifics. We computed two statistics, *d*′ and H, where *d*′ is signal strength (a higher value indicates that the mating signal from conspecifics is more readily detected), and H reflects an individual’s mating strategy. H ≍ 1.0 indicates unbiased decision making; H ≍ 0.0 indicates a bias towards mating with either a conspecific or heterospecific individual (i.e., a liberal strategy); and H > 1.0 indicates a bias towards rejection of mating with either a conspecific or heterospecific individual (i.e., a conservative strategy). *d*′ and H in response to signals (male mating attempt and female rejection behaviour) in each species were computed by using the dprime function of the neuropsychology library for the R software package (Makowski 2017). To visualize the decision-making performance in response to both male mating attempts and female rejection behaviour, we calculated the receiver operating characteristic (ROC) curve, which compares the sensitivity (the true positive rate, plotted on the *y*-axis) with the specificity (the false positive rate, plotted on the *x*-axis), for the signal detection results by using the ROCR package for R (Sing et al. 2005). Essentially, the closer an ROC curve is to the upper left corner, the better the decision-making accuracy, and the closer the curve is to the diagonal line of the panel (i.e., *y* = *x*), the more likely that the result is owing to chance alone (Carter et al. 2016). In addition, we used the DeLong method in the pROC package for R (Robin et al. 2011) to statistically compare the area under the ROC curve (AUC) between species in each experiment year.

### Comparative study

We performed a literature survey, using the ISI Web of Science (https://webofknowledge.com/) on 30 November 2017 and the key phrase “conspecific sperm precedence”, to identify congeneric pairs of animal species in which CSP had been detected in at least one of the pair. In addition, we screened the reference lists of two review papers for CSP (Howard 1999, Marshall et al. 2002) to locate additional pairs. We classified the geographic distributions and niches of each pair into one of four categories: (1) sympatry, geographical distribution of the two species largely overlaps with little if any niche separation in the sympatric area; (2) niche partitioning, geographical distributions of the two species overlap with niche partitioning at local scale (e.g., separation by food, habitat, or seasonality) especially at the reproductive stage; (3) parapatry, geographical distributions of the two species do not overlap but are adjacent with a narrow contact (hybridization) zone; or (4) allopatry, geographical distributions of the two species do not overlap and are not adjacent.

We excluded species with cosmopolitan, human-mediated distributions (e.g., *Drosophila simulans*, *Tribolium* flour beetles, and *Callosobruchus* bean weevils) from the analysis because their habitats and distributions in the natural environment are unclear. In total, we analysed 24 species pairs of marine invertebrates, terrestrial insects, and vertebrates.

## Results

### Mathematical model

#### Equilibria

We found dynamic population equilibria, designated by an asterisk (*), on (i) the *N*_X_-axis (i.e., 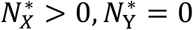), (ii) the 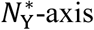 (i.e., 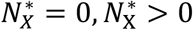), or (iii) in the interior (i.e., 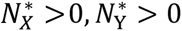). The boundary equilibria (as a result of competitive exclusion) are given by

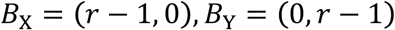

whereas the interior equilibrium did not have analytical formula (note that because we assume *r* > 1, boundary equilibria were always feasible).

#### Stability analyses

The stability conditions for the equilibria (species exclusion or coexistence) were determined from the eigenvalues of a Jacobi matrix around the focal equilibrium (more details are given in Appendix B). The necessary condition for a stable equilibrium resulting in extinction of one of the two species is given by:

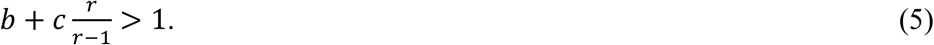

In particular, 4 = 1 (i.e., reproductive niches of the species overlap completely) necessarily leads to competitive exclusion given the parameter set for *p* and *q* used in our analysis (Fig. 2), even in the absence of interspecific resource competition (i.e., *b* = 0). See Appendix C for the numerical procedures for basins of attraction.

**Fig. 2.**
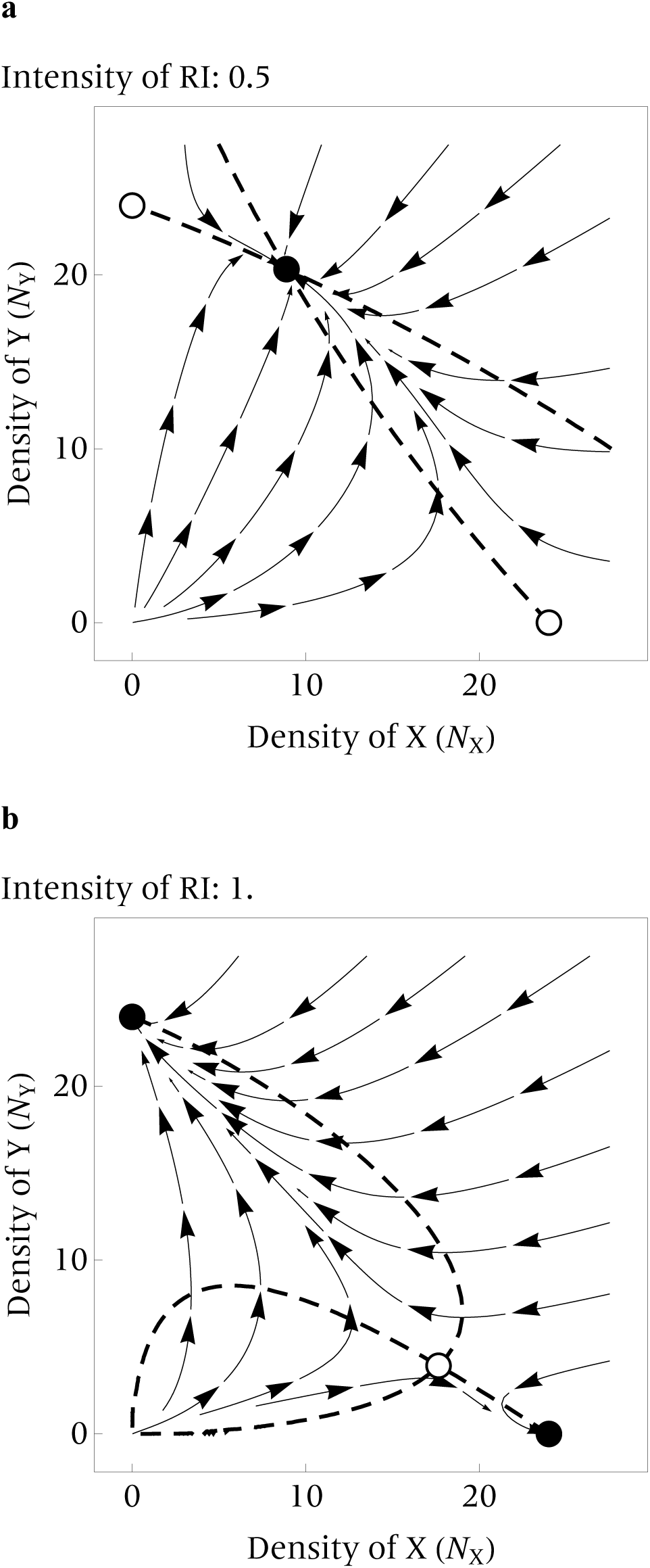
Phase portraits of the population dynamics according to the original, time-discrete dynamics and the approximated, time-continuous dynamics (i.e. ODE), for varying intensities of reproductive interference (RI; tuned by 4). (a) Co-existence is possible when RI intensity is weak (4 = 0.5). (b) Competitive exclusion occurs when RI intensity is very strong (4 = 1). Dotted curves, isoclines; arrows: approximated vector fields based on the ODE; open circles, unstable equilibria; and closed circles, stable equilibria. The procedure used to produce the figures is described in Appendix C. Probability parameter values: (*p*_XX_ = 0.4, *q*_XX_ =.4, *p*_XY_ = 0.8, *p*_YY_ = 0.8, *p*_YX_ = 0.4, *q*_YY_ = 0.8; other parameters, default values (see Table 1).

We note here that, if the two species are highly symmetric in terms of *p* and *q* values, then more outcomes become possible; in particular, species exclusion and coexistence states can be stable simultaneously (“bi-stable”), in agreement with Kishi and Nakazawa (2013) and Kyogoku and Sota (2017). Our particular intention here, however, is to explore the effects of asymmetry in mating behaviour (*p* and *q* values) on the community dynamics in our experimental system. For more details about the consequences of symmetric *p* and *q* values, see Appendix D. Also, it is possible to incorporate differences in the number of mating attempts in a given time period (i.e., mating activity) such that the encounter rate with an X- or Y-male can be biased towards either species relative to their frequency i the community (*f*_X_ and *f*_Y_); however, changes in the encounter rate did not change the results dramatically, although species exclusion became more likely (see Appendix D for more information).

### Experiment

Mating experience did not have a significant effect on the rate of copulation in either the 2014 or the 2015 experiment (Fig. SI 4, Table S1); therefore, virgin and non-virgin females were pooled in the following analyses. The copulation rate was higher in *H. axyridis* females than in *H yedoensis* females, especially in the 2014 experiments, although the difference was not statistically significant (Fig. 3, Table S2). In the 2015 experiment, *H. axyridis* was more likely to mate with conspecifics, whereas no such assortative mating pattern was observed in *H. yedoensis*; that is there was a significant interaction effect between female species and species identity of the mating partner (conspecific or heterospecific; Fig. 3, Table S2). In both the 2014 and 2015 experiments, *H. axyridis* males more frequently attempted to mate with conspecific females, whereas *H. yedoensis* males did not show a significant preference towards conspecific females (Table S3). *Harmonia axyridis* females were more likely than *H. yedoensis* females to refuse mating attempts by conspecific males, especially in the 2014 experiment (Table S4); however, both coercive mating and copulation failure occurred in both species following female rejection behaviour.

**Fig. 3.**
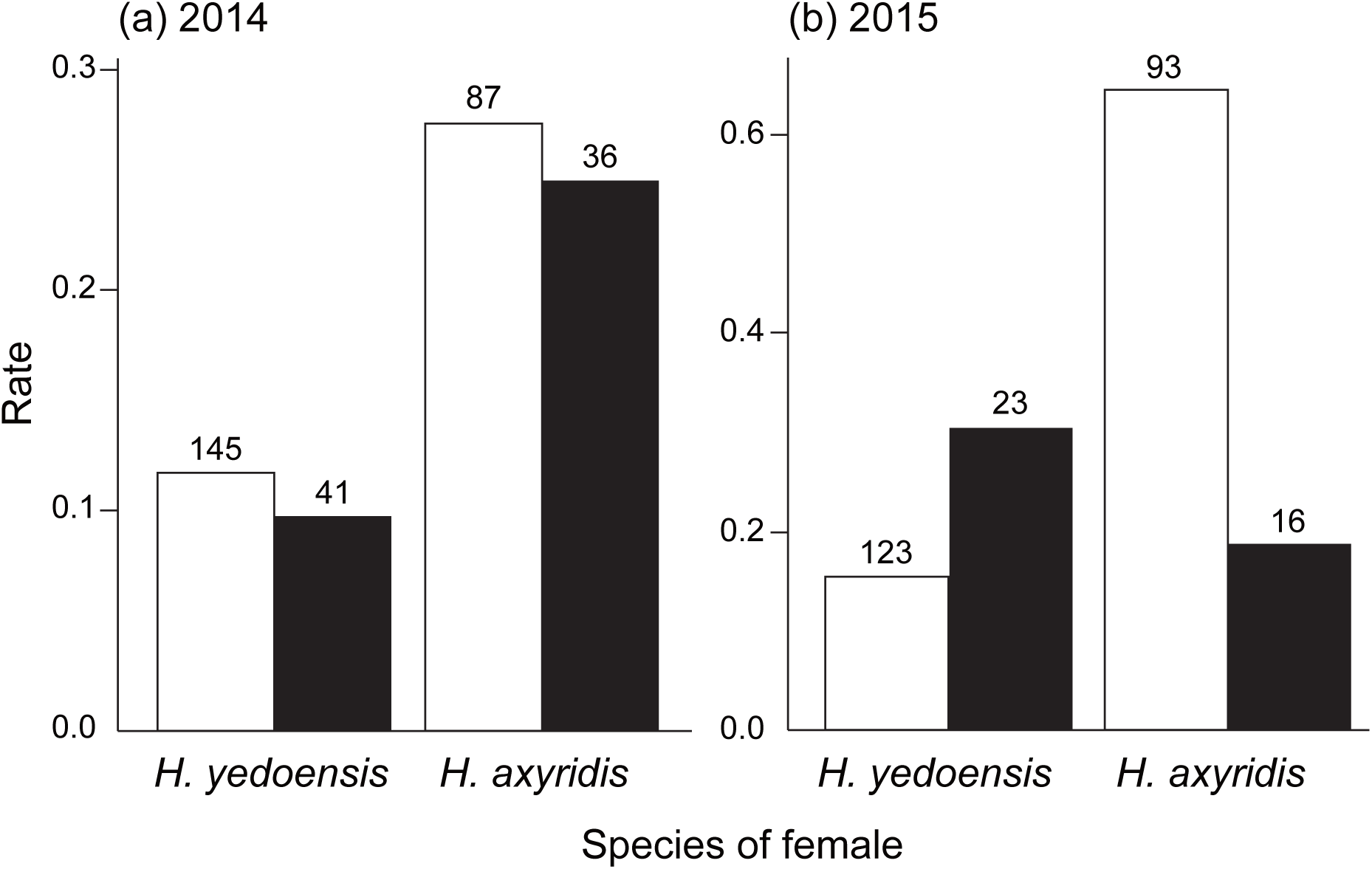
Mating rates with conspecific (white) and heterospecific (black) males in the (a) 2014 and (b) 2015 experiments. The number of individuals in each category is shown above each bar.

In the signal detection analysis results, *d*′ in response to male mating attempts was higher in *H. axyridis* than in *H. yedoensis* in both 2014 and 2015 (Table S5). Further, the AUC for male mating attempts was significantly higher in *H. axyridis* than in *H. yedoensis* in both 2014 and 2015 (Fig. 4, Table 2). By contrast, no consistent pattern in female rejection behaviour was detected between species or years in the signal detection analysis, probably in part because of the small sample size (Table S6). The AUC results for female rejection behaviour was also not significantly different between species in either experiment year (Fig. 4, Table 2). (c) Comparative study

**Fig. 4.**
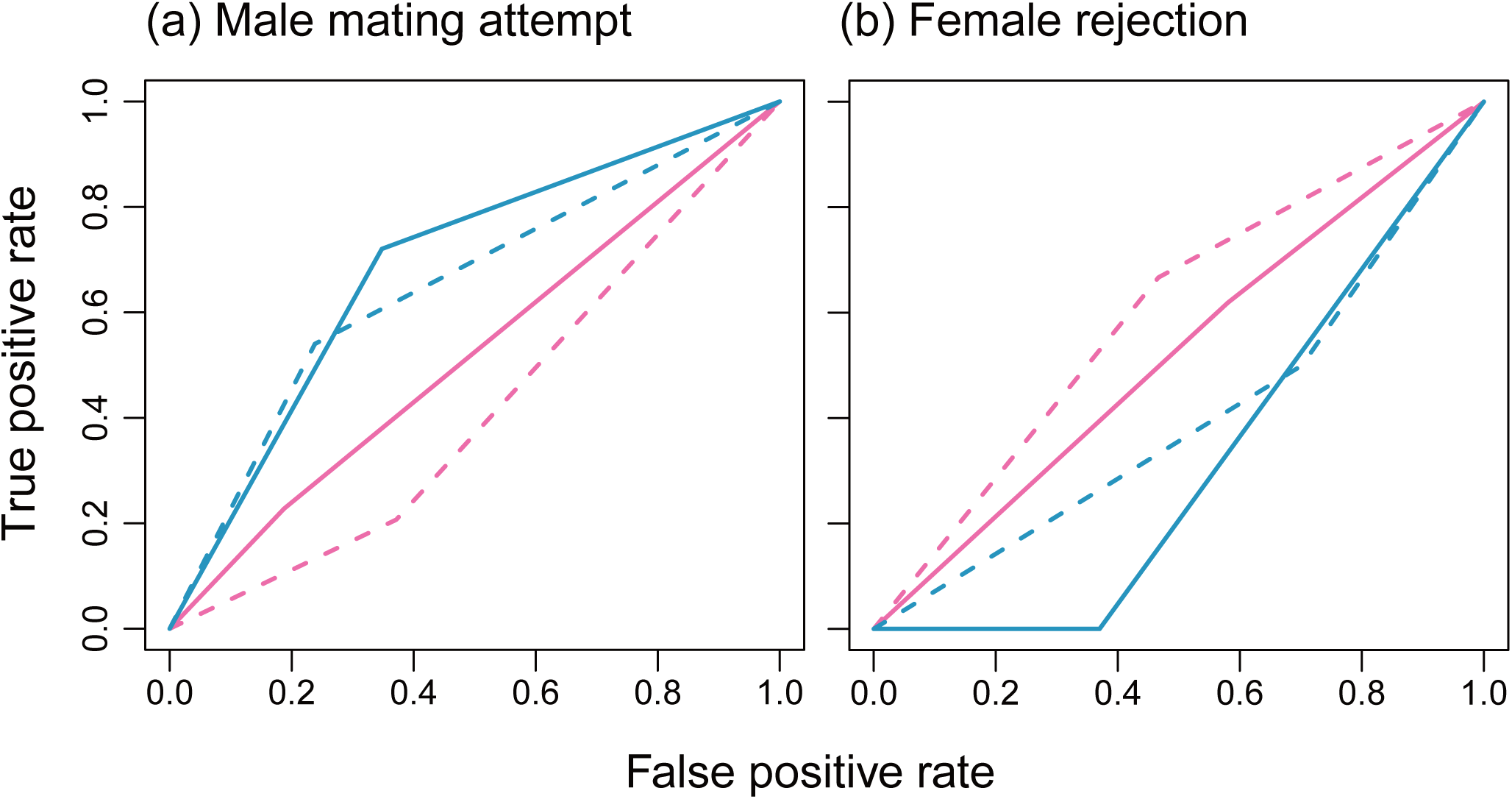
Receiver operating characteristic curves for (a) mating attempts by males and (b) rejection behaviour in females. In each panel, red and blue lines indicate *H. yedoensis* and *H. axyridis*, respectively, and dashed and solid lines indicate the 2014 and 2015 experiments, respectively.

**Table 2.**
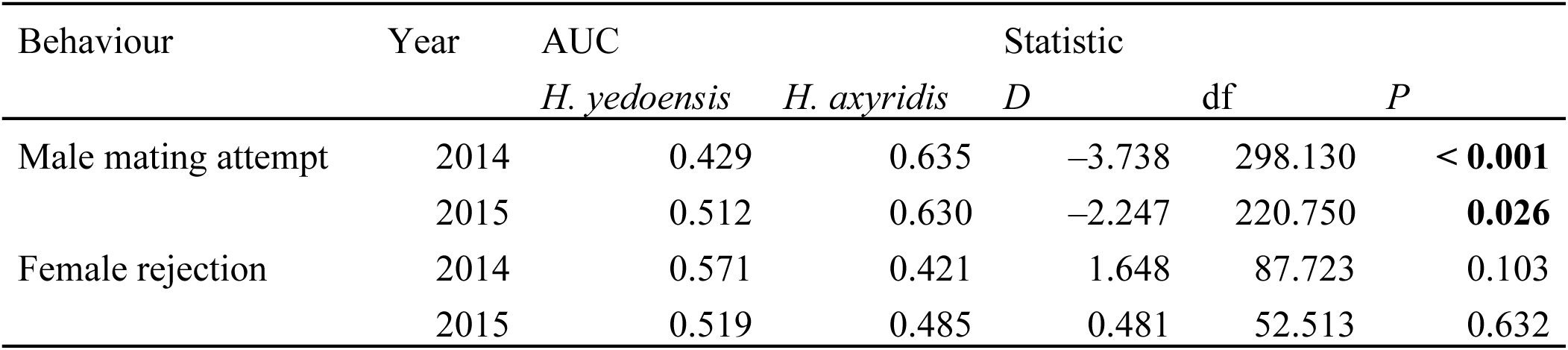
Comparison of the area under curve (AUC) values between species. Statistically significant results are shown in boldface.

| Behaviour | Year | AUC |  | Statistic |  |  |
| --- | --- | --- | --- | --- | --- | --- |
|  |  | *H. yedoensis* | *H. axyridis* | *D* | df | *P* |
| Male mating attempt | 2014 | 0.429 | 0.635 | −3.738 | 298.130 | < **0.001** |
|  | 2015 | 0.512 | 0.630 | −2.247 | 220.750 | **0.026** |
| Female rejection | 2014 | 0.571 | 0.421 | 1.648 | 87.723 | 0.103 |
|  | 2015 | 0.519 | 0.485 | 0.481 | 52.513 | 0.632 |

We found spatial separation at both local (niche partitioning) and regional scales (parapatry or allopatry) among species pairs exhibiting CSP, including in marine abalones, freshwater fishes, terrestrial insects, birds, and mice (Table 3). We observed parapatry mainly in Orthoptera (crickets and grasshoppers). We detected sympatry without apparent niche partitioning in 6 of 24 species pairs, especially in aquatic invertebrates such as mussels, starfishes, and sea urchins.

**Table 3.**
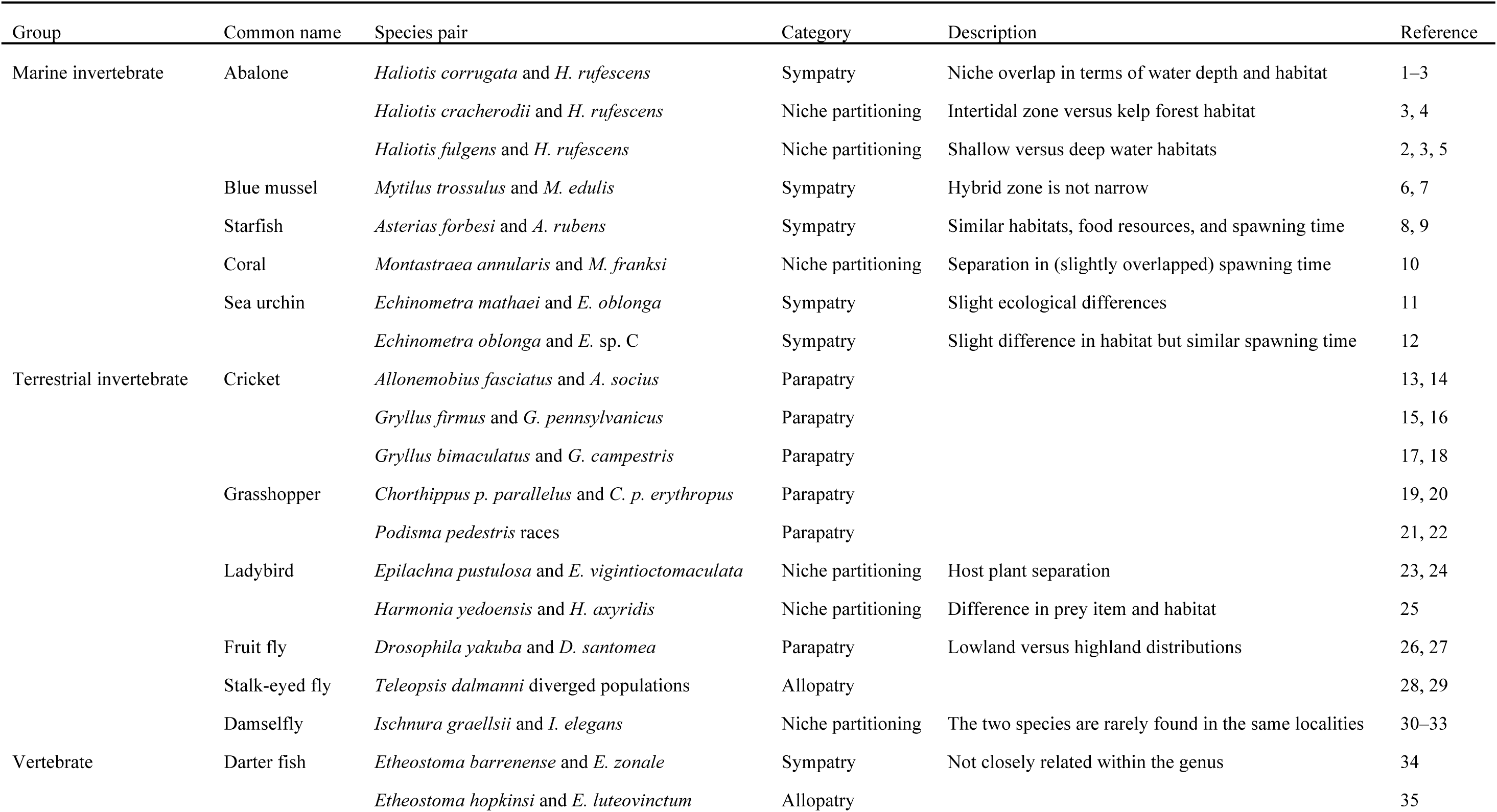
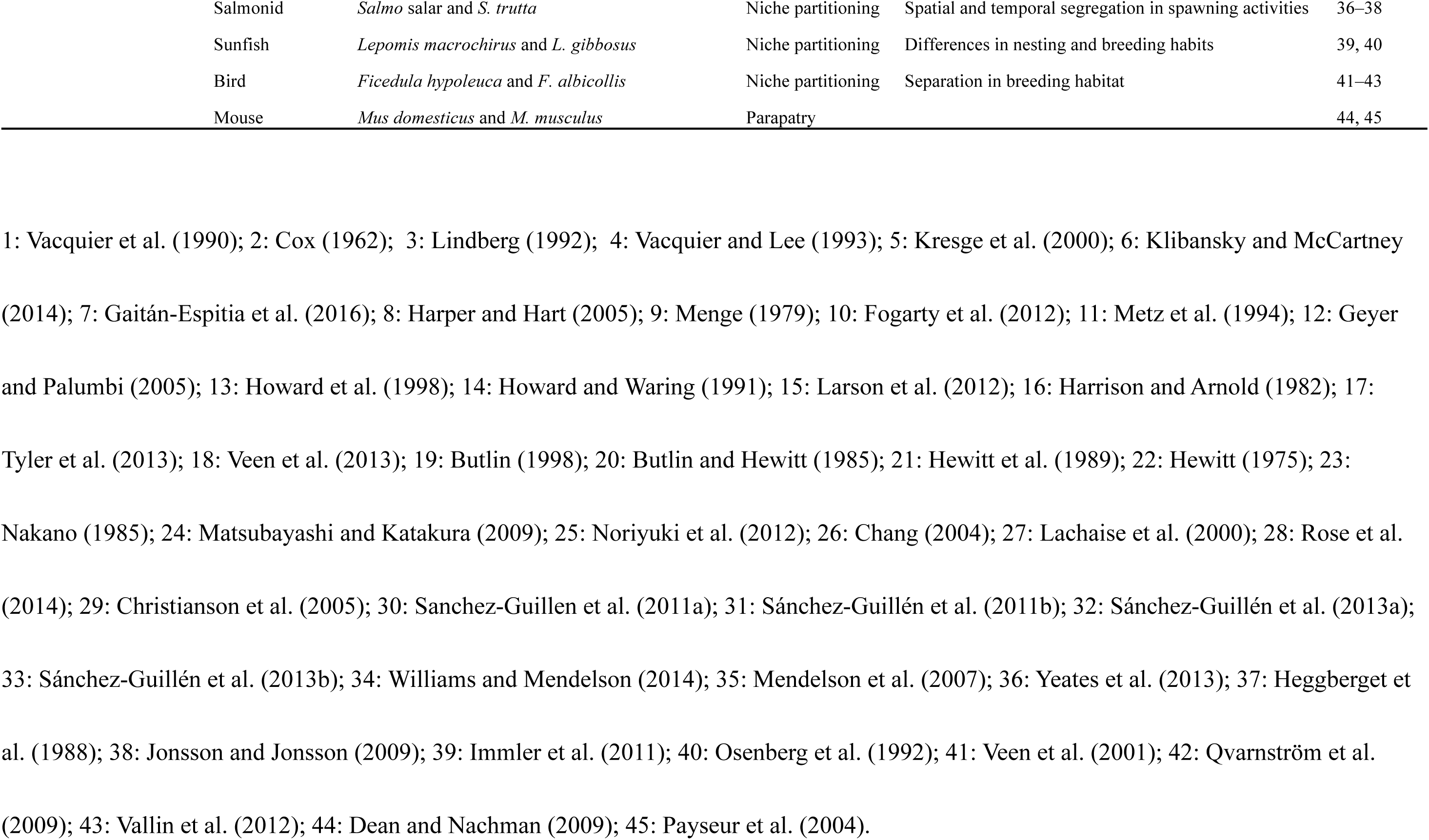
Summary of comparative study results.

## Discussion

Our results suggest that reproductive interference is likely to hamper stable species coexistence in a local patch even when the interacting species exhibit CSP. Our experimental results for two species in which CSP has been detected showed that the mating rate in a given period was higher in *H. axyridis* females than in *H. yedoensis* females (Fig. 3), and that *H. axyridis*, but not *H. yedoensis*, was more likely to copulate with a partner of its own species (Figs. 3 and 4). Our mathematical model indicated that these observed behavioural differences between these *Harmonia* species have a community-level consequence: namely, *H. yedoensis* becomes extinct in a local patch because of reproductive interference from *H. axyridis* (Figs. SI 2 and SI 3). Furthermore, our comparative study of species pairs exhibiting CSP demonstrated that parapatric distribution or niche partitioning, but not sympatric coexistence in the same habitat, can be maintained between two closely related species of a wide range of taxa, including both vertebrates and invertebrates living in either aquatic or terrestrial environments (Table 3). Taken together, these results lead us to conclude that CSP does not generally promote local coexistence between closely related species with overlapping reproductive niches.

Our experiment using *Harmonia* ladybirds, combined with our theoretical analysis, clarified the behavioural mechanisms of species exclusion. The rate of copulation was not significantly different between virgin and non-virgin females in the two *Harmonia* species (Fig. SI 4), suggesting that mating experience did not affect the reproductive success of individuals or the subsequent population dynamics in these species. However, the results of our signal detection analysis indicated that *H. axyridis* males easily distinguish and choose conspecific females over heterospecific females (Fig. 4, Table 2), whereas mating rates with conspecifics was low in *H. yedoensis* (Fig. 3). Our mathematical model demonstrated that, in the situations examined by our experiments, *H. axyridis* is likely to mate with a conspecific partner at least once before oviposition begins, whereas *H. yedoensis* females, even though they exhibit CSP, are incapable of producing viable offspring in the presence of *H. axyridis* males, with the result that *H. axyridis* is predicted to exclude *H. yedoensis* from the local patch (Fig. 2). This prediction is consistent with the niche partitioning observed in the field, where *H. axyridis* feeds on preferred prey items on various types of trees and *H. yedoensis* specializes in highly elusive prey on only pine trees. The pine habitat may function as a refuge for *H. yedoensis*, where it can avoid reproductive interference from *H. axyridis* (Noriyuki et al. 2012).

Our mathematical model highlighted the behavioural mechanisms that affect the asymmetry of reproductive interference and subsequent species exclusion. Although the classic theory of interspecific competition postulates that species exclusion occurs through exploitative competition for shared resources (Chesson 2000), our model results demonstrated that interference interactions during the reproductive stage hamper the coexistence of two species even when they demonstrate equal competitive strength for resources. In addition, our model results showed that slight differences in mating activity, mating preference, and remating acceptance determine which of two interacting species is superior with respect to reproductive interference (Figs. SI 2 and SI 3), whereas previous theoretical studies on reproductive interference did not fully take into account the consequences of behavioural processes on population dynamics and species’ fates (Yoshimura and Clark 1994, Kishi and Nakazawa 2013, Kyogoku and Sota 2017). In addition, we found that species exclusion is more likely to occur for a wide range of initial population densities of the two species when the intensity of reproductive interference is high (Fig. 2). This finding means that closely related species are unlikely to coexist in the same environment if they have similar mating signals or if they share a reproductive niche in space and time; as a result, niche partitioning or geographical segregation of the species is likely to occur.

Our comparative study found a separation of niche use or geographical distributions between species pairs with CSP in a range of taxa (Table 3). This finding suggests that CSP alone does not allow these species pairs to coexist in the same local environment. The pattern corresponds found in our comparative study is consistent with the prediction of our mathematical model that two interacting species are unlikely to coexist when *c* (intensity of reproductive interference) is high (Fig. 2). However, sympatric coexistence without apparent niche separation was also detected, especially in free-spawning marine invertebrates such as mussels, starfishes, and sea urchins (Table 3). There are several possible reasons that can account for the discrepancy between our model prediction and the actual pattern in nature in these cases. First, niche separation might actually exist, but, perhaps because of limited field survey data, it may not have been recognized. In fact, fine-scale differences in adult habitat and the timing of spawning have been detected in closely related marine invertebrate species (Lindberg 1992, Fogarty 2012). Therefore, it is possible that niche separation has actually occurred to mitigate the cost of reproductive interference in such species. Second, dispersal to new patches can allow overlapping niche use at a local scale even when two species engage in competitive interactions. Especially in marine sessile invertebrates that have high dispersal ability in the larval stage and a sedentary life style in the adult stage, source**–**sink dynamics (Mouquet and Loreau 2003) and stochastic processes (Paine and Levin 1981) likely promote local species coexistence. Third, in sessile animals, decision making at the pre-mating stage may not be important; females may be likely to accept sperm from conspecific as well as from heterospecific males, which means that CSP makes it possible for them to produce viable offspring. In this situation, therefore, CSP can indeed mitigate the cost of interspecific mating and thus promote species coexistence in the same niche. Clearly, it is important to incorporate life-history characteristics when considering the community-level consequences of behavioural decision making in animals.

By including plants, it would be possible to extend our model to more general scenarios of interacting species under imperfect species recognition. Reproductive interference occurs in flowering plants when the stigma receive heterospecific as well as conspecific pollen grains, for example when flowering phenology and pollinators overlap (Matsumoto et al. 2010, Runquist and Stanton 2013, Takakura 2013, Nishida et al. 2014). In some cases, however, conspecific pollen tubes preferentially grow and fertilize the ovules (Baldwin and Husband 2010, reviewed in Howard 1999). This phenomenon is called conspecific pollen precedence, and is considered a mechanism of reproductive isolation that prevents hybridization, and consequently, speciation in plants (Howard 1999). Therefore, it is suggested that conspecific pollen precedence in plants, similar to CSP in animals, can mitigate the cost of reproductive interference and lead to species coexistence in the same habitat.

Alternatively, as our model predicted, conspecific pollen precedence may be insufficient to allow interacting species to coexist in the same local environment. In fact, in three species of *Iris*, conspecific pollen precedence has been detected together with habitat differences (Carney et al. 1996; Emms et al. 1996), suggesting that reproductive interference destabilizes local coexistence of these species. In future, it would be interesting to examine whether our model is applicable to plant species by investigating reproductive success in species pairs exhibiting conspecific pollen precedence.

In conclusion, our study clarified the ecological significance of CSP by identifying conditions that lead to local species exclusion despite the presence of CSP. This finding is in contrast to those of previous studies of CSP, which have focused on its evolutionary significance, that is, speciation through post-mating pre-zygotic reproductive isolation.

Moreover, many CSP studies have not quantified pre-mating behaviours that can affect the reproductive success of females but have instead examined the functioning of CSP by focusing on post-mating, pre-zygotic mechanisms. Importantly, however, it has been documented that the overall costs of reproductive interference, including loss of mating opportunity and decreases in the oviposition rate due to male interference, can lead to the extinction of one of the interacting species even if interspecific mating and insemination does not occur (Kishi et al. 2009, Friberg et al. 2013, Carrasquilla and Lounibos 2015). Therefore, to understand individual reproductive success and community structure of closely related species, pre-mating behaviours should not be neglected.

## Acknowledgements

S.N. would like to thank staff members at the University of Tokyo Tanashi Forest for permission to collect insects. This study was supported grants from the Natural Environment Research Council (NERC) to R.I. (NE/K014617/1) and the Japan Society for the Promotion of Science to S.N (No. 26840137).

## Declarations

The authors declare no competing interest.

