## Supplementary Materials for "Reproductive interference hampers species coexistence despite conspecific sperm precedence"

(syllabarily ordered:) Ryosuke IRITANI [*†]& Suzuki NORIYUKI [‡§]

March 22, 2018

#### Contents

**A Reproductive output** **2**

**B Stability analysis** **3**

**C Numerical procedures for basins of attraction** **4**

**D General case** **6**

4.1. Effects of $p_{\mathrm{X|Y}}$ and $p_{\mathrm{Y|X}}$ . . . 6

4.2. Differential mating activity . . . 8

**E Summary of GLMM and signal detection indices** **10**

[*]Department of Integrative Biology, University of California, Berkeley, USA

[†]Biosciences, College of Life and Environmental Science, University of Exeter, Cornwall Campus, Penryn, Cornwall TR10 9EZ, the United Kingdom

[‡]Faculty of Agriculture and Marine Science, Kochi University, Japan

#### A Reproductive output

We assume that a female in state-$(i, j)$ is such that mated with a X-male $i$-times, and with a Y-male $j$-times. For instance, a state-$(0, 0)$ females are unmated. Let $\phi_{(i,j)}$ be the frequency of $(i, j)$-state X-females after the stage of mating.

The (2,0)-state is a set of females that mated with conspecifics twice, which occurs via: (i) rejecting males $k$-times successively (which occurs with probability $\xi_k$), (ii) encountering a X-male (which occurs with $f_X$), (iii) accepting him (which occurs with $p_{X|X}$), (iv) again encountering a X-male (which occurs with $f_X$), and (v) accepting him (which occurs with $q_{X|X}$). Bearing in mind the "loop" present in the diagram in the main text, we can show that:

$$\xi_k = \left( \underbrace{f_X}_{\text{encountering X-male}} \underbrace{(1 - p_{X|X})}_{\text{rejecting him}} + \underbrace{f_Y}_{\text{encountering Y-male}} \underbrace{(1 - p_{X|Y})}_{\text{rejecting him}} \right)^k = \left(1 - \overline{p_X}\right)^k, \tag{A-1}$$

where $\overline{p_X} := f_X p_{X|X} + f_Y p_{X|Y}$ represents the average acceptance rate of a unmated X-female. At the $(k + 1)$-st and $(k + 2)$-nd mating trials, she accepts the mating attempts from conspecifics, which occurs with probability $f_X p_{X|X} \cdot f_X q_{X|X}$. Hence the frequency of $(2, 0)$-females reads:

$$\phi_{(2,0)} = \sum_{k=0}^{\infty} \xi_k f_X p_{X|X} f_X q_{X|X} = \frac{p_{X|X} f_X^2 q_{X|X}}{\overline{p_X}} = \frac{p_{X|X} f_X^2 q_{X|X}}{f_X p_{X|X} + f_Y p_{X|Y}}, \tag{A-2}$$

following a basic result of the sum of geometric series.

Similarly, we get:

$$\begin{aligned} \phi_{(1,1)} &= \frac{f_X p_{X|X} f_Y q_{X|Y} + f_Y p_{X|Y} f_X q_{X|X}}{f_X p_{X|X} + f_Y p_{X|Y}}, \\ \phi_{(1,0)} &= \frac{f_X p_{X|X} f_X (1 - q_{X|X}) + f_X p_{X|X} f_Y (1 - q_{X|Y})}{f_X p_{X|X} + f_Y p_{X|Y}}, \\ \phi_{(0,1)} + \phi_{(0,2)} &= 1 - \left( \phi_{(2,0)} + \phi_{(1,1)} + \phi_{(1,0)} \right). \end{aligned} \tag{A-3}$$

Note that we are not required to explicitly evaluate the quantity on the final line, because the females in state $(0, 1)$ and $(0, 2)$ have not received conspecific sperms and thus have null fecundity.

From this, the expected reproductive output of a X-female per capita, $E_{\mathrm{X}}$, reads:

$$\begin{aligned} E_{\mathrm{X}} &= (1-c)r + cr\left(\phi_{(2,0)} + \phi_{(1,1)} + \phi_{(1,0)}\right) \\ &= (1-c)r + cr\frac{f_{\mathrm{X}}}{f_{\mathrm{X}}p_{\mathrm{X|X}} + f_{\mathrm{Y}}p_{\mathrm{X|Y}}}\left(p_{\mathrm{X|X}} + p_{\mathrm{X|Y}}q_{\mathrm{X|X}}f_{\mathrm{Y}}\right), \end{aligned} \tag{A-4}$$

where $r$ represents the total number of offspring surviving to maturation, per capita. Switching the subscripts X and Y on each symbol supplies the corresponding quantities for a Y-female.

Altoghether, the community dynamics is described as:

$$\begin{aligned} N_{\mathrm{X}}(t+1) &= \frac{N_{\mathrm{X}}(t)E_{\mathrm{X}}(t)}{1 + vN_{\mathrm{X}}(t) + bvN_{\mathrm{Y}}(t)} =: W_{\mathrm{X}}\left(N_{\mathrm{X}}(t), N_{\mathrm{Y}}(t)\right), \\ N_{\mathrm{Y}}(t+1) &= \frac{N_{\mathrm{Y}}(t)E_{\mathrm{Y}}(t)}{1 + bvN_{\mathrm{X}}(t) + vN_{\mathrm{Y}}(t)} =: W_{\mathrm{Y}}\left(N_{\mathrm{X}}(t), N_{\mathrm{Y}}(t)\right). \end{aligned} \tag{A-5}$$

We can without loss of generality assume $v = 1$ by subsuming $v$ with the densities. Indeed, suppose that $v = 1$; for any $\hat{v} > 0$, we can get:

$$\begin{aligned} \hat{v}N_{\mathrm{X}}(t+1) &= W_{\mathrm{X}}\left(\hat{v}N_{\mathrm{X}}(t), \hat{v}N_{\mathrm{Y}}(t)\right), \\ \hat{v}N_{\mathrm{Y}}(t+1) &= W_{\mathrm{Y}}\left(\hat{v}N_{\mathrm{X}}(t), \hat{v}N_{\mathrm{Y}}(t)\right), \end{aligned} \tag{A-6}$$

so that we can transform the variable $N_i(t)$ to $n_i(t) = \hat{v}N_i(t)$, while $E_i$ is unchanged with such a transformation (for $i =$X, Y).

#### B Stability analysis

Here we will concentrate on the original, discrete-time dynamics for the stability. Readers may wonder if and when the discrete-time and continuous-time agree with each other on the stability, but the small changes in density values over discrete-time would guarantee that these approaches yield the same stability conditions (Otto & Day 2007, Chapter 2).

At equilibrium,

$$\begin{aligned} \frac{N_{\mathrm{X}}E_{\mathrm{X}}}{1 + N_{\mathrm{X}} + bN_{\mathrm{Y}}} &= N_{\mathrm{X}}, \\ \frac{N_{\mathrm{Y}}E_{\mathrm{Y}}}{1 + N_{\mathrm{Y}} + bN_{\mathrm{Y}}} &= N_{\mathrm{Y}}. \end{aligned} \tag{B-1}$$

$N_X = N_Y = 0$ is necessarily a trivial, unstable equilibrium, given that $r > 1$ (Ackleh & Salceanu 2014).

##### *Species exclusion*

It is easy to show that $B_X^* = (r-1, 0)$ and $B_Y^* = (0, r-1)$ are the boundary equilibria. $B_X^*$ is locally stable if:

$$\left| \left. \frac{\partial W_X(N_X, N_Y)}{\partial N_X} \right|_{B_X^*} \right| < 1, \qquad \left. \frac{\partial W_Y(N_X, N_Y)}{\partial N_Y} \right|_{B_X^*} < 1, \tag{B-2}$$

which finally translates to:

$$r > 1, \qquad b + c\frac{r}{r-1} < 1. \tag{B-3}$$

Similar computation yields the local stability condition of $B_Y^*$, which reads:

$$r > 1, \qquad b + c\frac{r}{r-1} < 1. \tag{B-4}$$

Therefore, the boundary equilibrium $B_X^*$ is stable if and only if $B_Y^*$ is stable.

##### *Coexistence*

Coexistence state $C^* = (N_X^*, N_Y^*)$ occurs when the absolute value of the dominant eigenvalue of the Jacobi matrix ($\mathcal{J}$) around $C^*$ is smaller than unity. It is unfortunately impossible to derive the analytical expression for the stability condition, and hence we carried out numerical investigation for the stability.

#### C Numerical procedures for basins of attraction

Using Mathematica (Wolfram Research 2016), we visualized the dynamics in the main text. We outline the minimal procedure. First, we used the `StreamPlot` function to produce the vector fields. Second, we used the `RecurrenceTable` function to iterate the original, discrete system over 70 generations (which we found is enough for the convergence). Here, to illustrate the basin of attraction, we produced pairs of initial values for $(x_0, y_0)$ uniformly over the plotting region (which is $1.5r \times 1.5r$ with fineness $\delta$), and checked what initial values recursively converge to which. Since we had already known that the species exclusion occurs for a large $c$, we put $\delta = 0.70$ if $c > 0.8$ or otherwise $\delta = 2$, using the `If` function. Note that as $\delta$ decreases the graphics figures become heavier. We then extracted $N_{\mathrm{X}}(70)$ and $N_{\mathrm{Y}}(70)$ starting from each initial condition produced above, and measured the distance between $\left(N_{\mathrm{X}}(70), N_{\mathrm{Y}}(70)\right)$ and $B^*_{\mathrm{X}} = (r-1, 0)$ (or $B^*_{\mathrm{Y}} = (0, r-1)$, respectively), using the `EucledeanDistance` function which we here denote as $d_{\mathrm{X}}$ (or $d_{\mathrm{Y}}$, respectively). If $d_{\mathrm{X}}$ (or $d_{\mathrm{Y}}$) is smaller than a very small threshold value $\varepsilon = 1/1000$, then we issue the option `PlotStyle->Blue` (or `PlotStyle->Red`, respectively), or otherwise `PlotStyle->Gray` for the function `ListPlot`, using the `If` function.

Finally, to illustrate the stability of the equilibria (open and closed circles), we numerically evaluated the dominant eigenvalue of the system around the equilibria. To do so, we counted the number of equilibria using the `Length` function, where the possible solutions are produced by the `NSolve` function applied to Eqn (A-5) (with a restricting condition $N_{\mathrm{X}} \geq 0$ and $N_{\mathrm{Y}} \geq 0$). We then produced a list of equilibria by using the `Table` function to obtain the dominant eigenvalues of the corresponding equilibria. We assessed if the dominant eigenvalue is larger than unity or not, and issued the option `EdgeForm` of the `Graphics` function such that unstable (or stable) points are filled in white (or black; FigSI 1).

We changed the colors and/or opacity depending on the outcomes (to enhance the readability), but we do not detail them here.

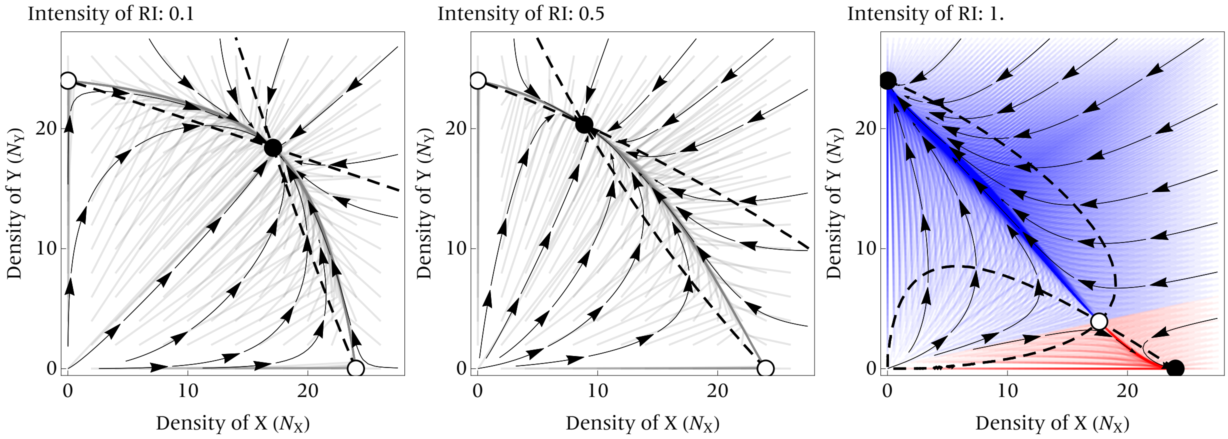


FigSI 1: Isoclines and phase portraits for the community dynamics. Starting with gray points (left and middle), the dynamics will converge to coexistence state. Starting with red or blue points (right), the dynamics converge will converge to boundary states. Initial points are generated randomly within the rectangles. Parameters are set at default values while $c$ is varied (0.1, 0.5 or 1).

### D General case

##### *4.1. Effects of $p_{X|Y}$ and $p_{Y|X}$*

From behavioral observation data, we used highly asymmetric parameter sets:

$$p_{X|X} = q_{X|X} = 0.4,\; p_{X|Y} = 0.8; \tag{D-1}$$

$$p_{Y|Y} = q_{Y|Y} = 0.8,\; p_{Y|X} = 0.4. \tag{D-2}$$

However, when these values are relatively symmetric, the community dynamics consequences are more divergent. In particular, with symmetry in $p$'s and $q$'s, there would be a bi-stability (Kuno 1992; Kishi & Nakazawa 2013; Kyogoku & Sota 2017).

For the completeness, we here illustrate the effects of symmetricity in $p$'s and $q$'s on the community dynamics by plotting phase portraits. To this end, it is useful to reduce the parameters by the following equations:

$$\begin{aligned} E_X &= (1-c)r + cr\frac{f_X}{f_X + \frac{p_{X|Y}}{p_{X|X}} f_Y}\left(1 + \frac{p_{X|Y}}{p_{X|X}} q_{X|X} f_Y\right), \\ E_Y &= (1-c)r + cr\frac{f_Y}{f_Y + \frac{p_{Y|X}}{p_{Y|Y}} f_X}\left(1 + \frac{p_{Y|X}}{p_{Y|Y}} q_{Y|Y} f_X\right) \end{aligned} \tag{D-3}$$

so that we can subsume the fractions into compound parameters:

$$\frac{p_{X|Y}}{p_{X|X}} = \pi_{X|Y}, \frac{p_{Y|X}}{p_{Y|Y}} = \pi_{Y|X} \tag{D-4}$$

that represent relative acceptance rate of a female for heterospecific males over conspecific males. To look at the effect of acceptance rates of females ($\pi_{X|Y}$ and $\pi_{Y|X}$), we fix $q := q_{X|X} = q_{Y|Y}$ at a particular value from 0, 0.5, to 1, and vary $\pi$'s (FigSI 2).

As we have found already, the stability for boundary equilibria ($B^*_X, B^*_Y$) is independent of $\pi$'s and $q$, while that for interior equilibria depends on $\pi$'s and $q$. As such, varying $\pi$'s and $q$ can generate the bistability of species exclusion and coexistence.

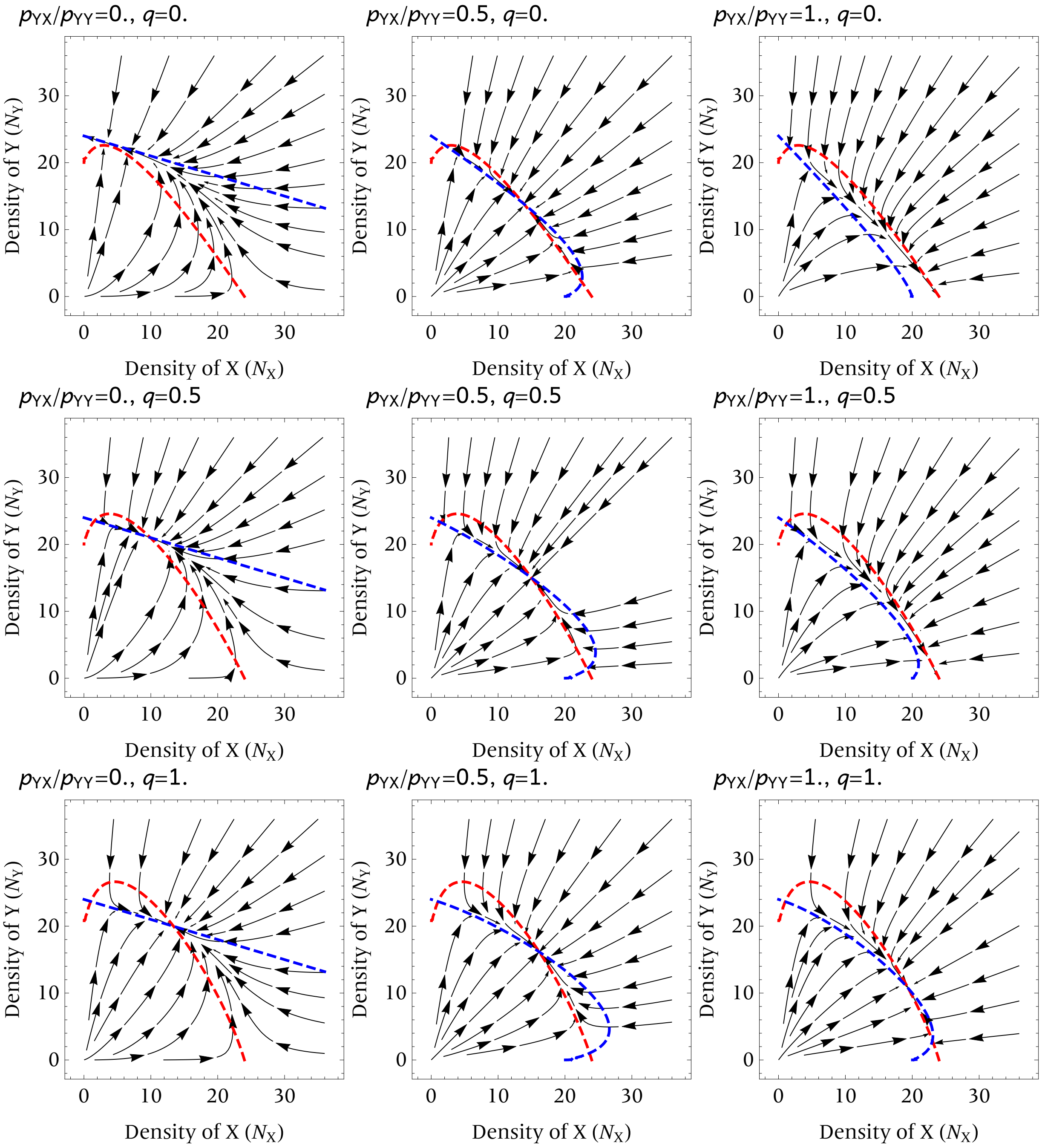


FigSI 2: Phase portraits, with $\pi_{\mathrm{Y|X}} = 0, 0.5, 1$ (from left to right) and $q = 0, 0.5, 1$ (from top to bottom), given $\pi_{\mathrm{Y|X}} = 0.5$ (fixed). From these panels, we can see that bistabitliy can occur. Red curves: isoclines for X; Blue curves: isoclines for Y.

##### *4.2. Differential mating activity*

The rate of mating attempt in a given time (hereafter mating activity) can differ remarkably between species (see Fig. 3 in the main text). To account for this, we define asymmetric frequencies $g_{\mathrm{X}}$ and $g_{\mathrm{Y}}$ by:

$$g_{\mathrm{X}} = \frac{N_{\mathrm{X}}}{N_{\mathrm{X}} + \alpha_{\mathrm{Y}} N_{\mathrm{Y}}} = 1 - g_{\mathrm{Y}}, \tag{D-5}$$

where $\alpha_{\mathrm{Y}}$ represents the relative mating activity of a Y-male over X-male ($\alpha_{\mathrm{Y}} > 0$). Higher $\alpha_{\mathrm{Y}}$ indicates a higher mating activity of Y-males; $\alpha_{\mathrm{Y}} = 1$ implies the even level of mating activity.

We replaced $f$'s by $g$'s:

$$\begin{aligned} E_{\mathrm{X}} &= (1-c)r + cr\frac{g_{\mathrm{X}}}{g_{\mathrm{X}} + \frac{p_{\mathrm{X|Y}}}{p_{\mathrm{X|X}}} g_{\mathrm{Y}}} \left(1 + \frac{p_{\mathrm{X|Y}}}{p_{\mathrm{X|X}}} q_{\mathrm{X|X}} g_{\mathrm{Y}}\right), \\ E_{\mathrm{Y}} &= (1-c)r + cr\frac{g_{\mathrm{Y}}}{g_{\mathrm{Y}} + \frac{p_{\mathrm{Y|X}}}{p_{\mathrm{Y|Y}}} g_{\mathrm{X}}} \left(1 + \frac{p_{\mathrm{Y|X}}}{p_{\mathrm{Y|Y}}} q_{\mathrm{Y|Y}} g_{\mathrm{X}}\right), \end{aligned} \tag{D-6}$$

and found that increasing $\alpha_{\mathrm{Y}}$ leads to higher possibility of the extinction of X; this is a natural result because mating activity can lead to higher conspecific mating-chance for X-females.

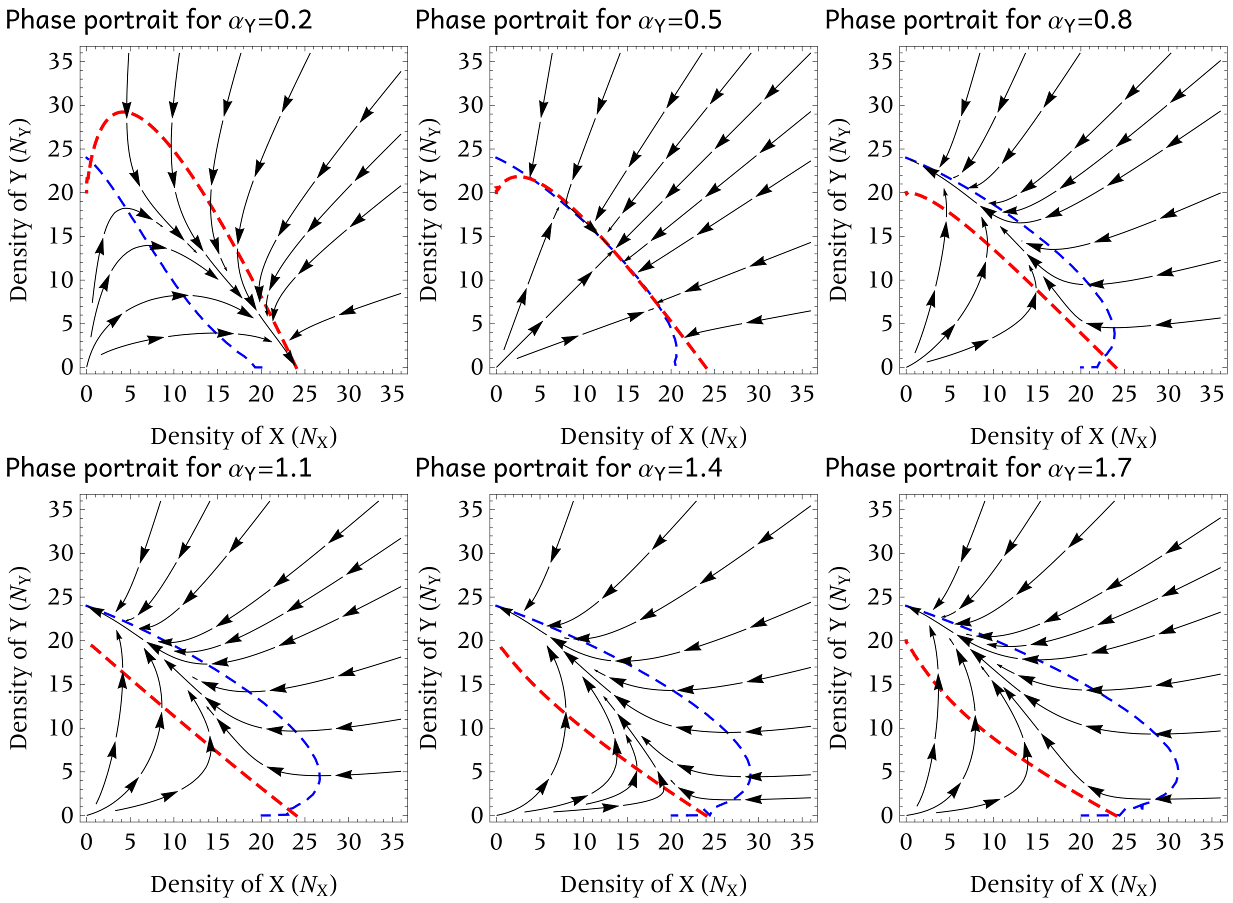


FigSI 3: Isoclines and phase portraits for the community dynamics. Red curves: isoclines for X; Blue curves: isoclines for Y. Higher $\alpha_Y$ leads to lower possibility of the persistence of X. Default parameter values are used.

#### E Summary of GLMM and signal detection indices

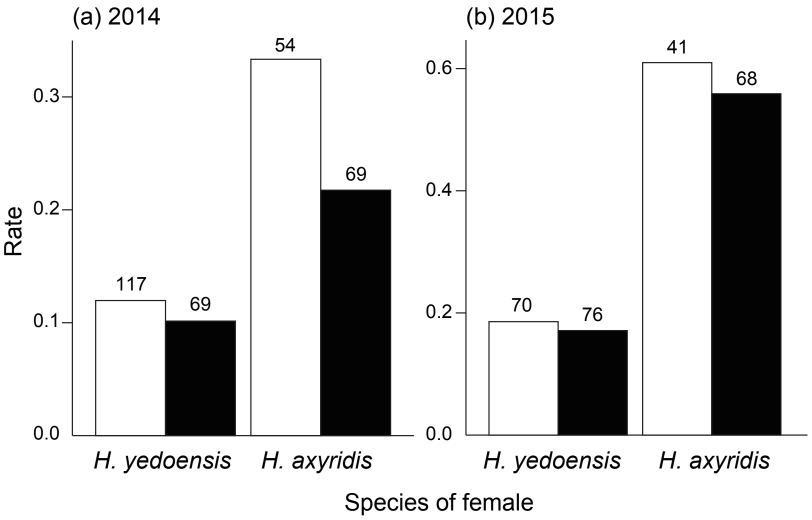


FigSI 4: Mating rate of unmated (white) and mated (black) females in (a) 2014 and (b) 2015 experiment. The sample size of each category is shown above each bar.

**Table S1.** Summary of GLMM of the mating experience and other factors affecting the mating rate in 2014 and 2015 experiment. Significant results are shown in bold.

| Year | Fixed effect | Estimate | SE | *z* value | *P* |
|---|---|---|---|---|---|
| 2014 | Intercept | 0.933 | 2.064 | 0.452 | 0.651 |
| | Mating experience | 0.080 | 0.521 | 0.153 | 0.878 |
| | Female species | 0.231 | 0.638 | 0.363 | 0.717 |
| | Mating experience * female species | 0.893 | 0.778 | 1.147 | 0.251 |
| | Female body length | –0.329 | 0.121 | –2.709 | **0.007** |
| | Male body length | –0.150 | 0.252 | –0.595 | 0.552 |
| | Female elytra colour | –0.041 | 0.449 | –0.090 | 0.928 |
| | Male elytra colour | 1.156 | 0.702 | 1.646 | 0.100 |
| | Female age | –0.031 | 0.013 | –2.341 | **0.019** |
| | Male age | 0.013 | 0.016 | 0.787 | 0.431 |
| 2015 | Intercept | –4.255 | 6.902 | –0.616 | 0.538 |
| | Mating experience | 0.440 | 0.718 | 0.612 | 0.541 |
| | Female species | 2.411 | 0.797 | 3.024 | **0.003** |
| | Mating experience * female species | 0.088 | 0.904 | 0.097 | 0.922 |
| | Female body length | –0.632 | 0.509 | –1.240 | 0.215 |
| | Male body length | 1.039 | 1.016 | 1.023 | 0.307 |
| | Female elytra colour | –0.241 | 0.550 | –0.437 | 0.662 |
| | Male elytra colour | –0.801 | 1.189 | –0.673 | 0.501 |
| | Female age | 0.063 | 0.050 | 1.257 | 0.209 |
| | Male age | –0.041 | 0.053 | –0.779 | 0.436 |

**Table S2.** Summary of GLMM of the intra– versus interspecific mating and other factors affecting the mating rate in 2014 and 2015 experiment. Significant results are shown in bold.

| Year | Fixed effect | Estimate | SE | *z* value | *P* |
|---|---|---|---|---|---|
| 2014 | Intercept | 0.249 | 2.068 | 0.120 | 0.904 |
| | Intraspecific mating | 0.487 | 0.670 | 0.726 | 0.468 |
| | Female species | 1.039 | 0.736 | 1.412 | 0.158 |
| | Intraspecific mating * female species | –0.425 | 0.969 | –0.438 | 0.661 |
| | Female body length | –0.319 | 0.120 | –2.656 | **0.008** |
| | Male body length | –0.158 | 0.264 | –0.597 | 0.550 |
| | Female elytra colour | 0.000 | 0.447 | 0.000 | 1.000 |
| | Male elytra colour | 1.146 | 0.693 | 1.653 | 0.098 |
| | Female age | –0.029 | 0.013 | –2.266 | **0.023** |
| | Male age | 0.016 | 0.016 | 1.024 | 0.306 |
| 2015 | Intercept | –2.534 | 6.832 | –0.371 | 0.711 |
| | Intraspecific mating | –2.824 | 1.505 | –1.876 | 0.061 |
| | Female species | –1.576 | 1.612 | –0.978 | 0.328 |
| | Intraspecific mating * female species | 6.606 | 2.895 | 2.282 | **0.023** |
| | Female body length | –0.598 | 0.504 | –1.187 | 0.235 |
| | Male body length | 1.012 | 1.019 | 0.993 | 0.321 |
| | Female elytra colour | –0.245 | 0.559 | –0.438 | 0.661 |
| | Male elytra colour | –0.973 | 1.213 | –0.802 | 0.423 |
| | Female age | 0.038 | 0.051 | 0.738 | 0.461 |
| | Male age | –0.014 | 0.054 | –0.260 | 0.795 |

**Table S3.** Summary of GLMM of the intra– versus interspecific mating and other factors affecting the male mating attempt in 2014 and 2015 experiment. Significant results are shown in bold.

| Year | Fixed effect | Estimate | SE | *z* value | *P* |
|---|---|---|---|---|---|
| 2014 | Intercept | –1.584 | 2.071 | –0.765 | 0.444 |
| | Intraspecific mating | –0.491 | 0.468 | –1.048 | 0.295 |
| | Male species | –0.451 | 0.644 | –0.701 | 0.484 |
| | Intraspecific mating * male species | 1.763 | 0.708 | 2.489 | **0.013** |
| | Female body length | –0.085 | 0.112 | –0.760 | 0.447 |
| | Male body length | 0.081 | 0.266 | 0.305 | 0.760 |
| | Female elytra colour | 0.567 | 0.432 | 1.313 | 0.189 |
| | Male elytra colour | 0.701 | 0.521 | 1.347 | 0.178 |
| | Female age | –0.011 | 0.009 | –1.131 | 0.258 |
| | Male age | 0.006 | 0.014 | 0.445 | 0.657 |
| 2015 | Intercept | –2.376 | 7.190 | –0.330 | 0.741 |
| | Intraspecific mating | –0.506 | 0.957 | –0.529 | 0.597 |
| | Male species | 1.957 | 1.771 | 1.105 | 0.269 |
| | Intraspecific mating * male species | 3.309 | 1.253 | 2.640 | **0.008** |
| | Female body length | –0.156 | 0.518 | –0.302 | 0.763 |
| | Male body length | 0.453 | 1.030 | 0.440 | 0.660 |
| | Female elytra colour | –0.742 | 0.576 | –1.287 | 0.198 |
| | Male elytra colour | –1.415 | 1.273 | –1.111 | 0.266 |
| | Female age | 0.053 | 0.054 | 0.984 | 0.325 |
| | Male age | –0.039 | 0.056 | –0.689 | 0.491 |

**Table S4.** Summary of GLMM of the intra– versus interspecific mating and other factors affecting the female rejection behaviour in 2014 and 2015 experiment. Significant results are shown in bold.

| Year | Fixed effect | Estimate | SE | *z* value | *P* |
|---|---|---|---|---|---|
| 2014 | Intercept | –6.204 | 4.421 | –1.403 | 0.161 |
| | Intraspecific mating | –1.059 | 0.878 | –1.207 | 0.228 |
| | Female species | –1.182 | 0.960 | –1.230 | 0.219 |
| | Intraspecific mating * female species | 2.351 | 1.145 | 2.054 | **0.040** |
| | Female body length | –0.067 | 0.165 | –0.408 | 0.683 |
| | Male body length | 1.195 | 0.683 | 1.751 | 0.080 |
| | Female elytra colour | 0.856 | 0.706 | 1.212 | 0.226 |
| | Male elytra colour | –0.312 | 0.894 | –0.349 | 0.727 |
| | Female age | 0.012 | 0.015 | 0.808 | 0.419 |
| | Male age | –0.015 | 0.022 | –0.706 | 0.480 |
| 2015 | Intercept | –3.722 | 6.102 | –0.610 | 0.542 |
| | Intraspecific mating | –0.126 | 0.859 | –0.147 | 0.883 |
| | Female species | –18.103 | 193.521 | –0.094 | 0.925 |
| | Intraspecific mating * female species | 17.025 | 193.520 | 0.088 | 0.930 |
| | Female body length | –0.082 | 0.627 | –0.130 | 0.896 |
| | Male body length | 0.467 | 0.674 | 0.692 | 0.489 |
| | Female elytra colour | 0.166 | 0.748 | 0.221 | 0.825 |
| | Male elytra colour | 1.495 | 1.084 | 1.379 | 0.168 |
| | Female age | 0.006 | 0.037 | 0.159 | 0.874 |
| | Male age | 0.003 | 0.040 | 0.076 | 0.940 |

**Table S5**. Results for the signal detection theory indicies.

| Behaviour | Year | Species | $d'$ | $\beta$ |
|---|---|---|---|---|
| Male mating attempt | 2014 | *H. yedoensis* | –0.480 | 0.760 |
| | | *H. axyridis* | 0.794 | 1.274 |
| | 2015 | *H. yedoensis* | 0.099 | 1.081 |
| | | *H. axyridis* | 0.960 | 0.921 |
| Female rejection | 2014 | *H. yedoensis* | 0.490 | 1.095 |
| | | *H. axyridis* | –0.487 | 0.871 |
| | 2015 | *H. yedoensis* | 0.098 | 1.026 |
| | | *H. axyridis* | –0.441 | 1.271 |

**Table S6**. The number of behavioural responses to correct signal (conspecific individuals) and false signal (heterospecific individuals) in mating attempt of males and rejection behaviour of females. Hit and Correct Rejection indicates right response to correct and false signals, respectively. Miss and False Alarm indicates wrong response to correct and false signals, respectively.

| Behaviour | Year | Species | Correct signal | | False signal | |
|---|---|---|---|---|---|---|
| | | | # Hit | # Miss | # False Alarm | # Correct Rejection |
| Male mating attempt | 2014 | *H. yedoensis* | 30 | 115 | 13 | 22 |
| | | *H. axyridis* | 47 | 40 | 10 | 32 |
| | 2015 | *H. yedoensis* | 28 | 95 | 3 | 13 |
| | | *H. axyridis* | 67 | 26 | 8 | 15 |
| Female rejection | 2014 | *H. yedoensis* | 16 | 14 | 3 | 6 |
| | | *H. axyridis* | 14 | 33 | 7 | 7 |
| | 2015 | *H. yedoensis* | 13 | 18 | 8 | 13 |
| | | *H. axyridis* | 97 | 57 | 3 | 0 |
